# 3D Bioprinting of Stem Cell-Derived Central Nervous System Cells Enables Astrocyte Growth, Vasculogenesis and Enhances Neural Differentiation/Function

**DOI:** 10.1101/2022.11.13.516338

**Authors:** Michael A. Sullivan, Samuel D. Lane, Alexander Volkerling, Martin Engel, Eryn L. Werry, Michael Kassiou

**Affiliations:** School of Medical Sciences, The Faculty of Medicine and Health, The University of Sydney, Sydney, Australia; School of Chemistry, The Faculty of Science, The University of Sydney, Sydney, Australia; Inventia Life Science Operations Pty Ltd, Alexandria, NSW 2015, Australia; Central Clinical School, Faculty of Medicine and Health, The University of Sydney

**Author notes:** Co-corresponding author: Michael Kassiou, **Email**. Co-corresponding author: Eryn Werry, **Email**.

**Keywords:** biomaterials, hydrogel, poly(ethylene glycol), 3D, bioprinting, iPSC, CNS

## Abstract

Current research tools for pre-clinical drug development such as rodent models and 2D immortalised monocultures have failed to serve as effective translational models for human CNS disorders. Recent advancements in the development of iPSCs and 3D culturing can improve the *in vivo-relevance* of pre-clinical models, while generating 3D cultures though novel bioprinting technologies can offer increased scalability and replicability. As such, there is a need to develop platforms that combine iPSC-derived cells with 3D bioprinting to produce scalable, tunable and biomimetic cultures for preclinical drug discovery applications. We report a biocompatible PEG-based matrix which incorporates RGD and YIGSR peptide motifs and full length collagen IV at a stiffness similar to the human brain (1.5 kPa). Using a high-throughput commercial bioprinter we report the viable culture and morphological development of iPSC-derived astrocytes, brain microvascular endothelial cells, neural progenitors and neurons in our novel matrix. We also show that this system supports endothelial vasculogenesis and enhances neural differentiation and spontaneous activity. This platform forms a foundation for more complex, multicellular models to facilitate high-throughput translational drug discovery for CNS disorders.

## 1. Introduction

Central nervous system (CNS) disorders are the leading cause of disability and second leading cause of death worldwide (Group, 2017). Understanding how cells function and communicate in the CNS is critical to tackling this issue, however the disordered human CNS consists of uniquely complex pathologies, which are prohibitively invasive to study at close proximity (Drummond & Wisniewski, 2017). For this reason, the ability of our *in vivo* and *in vitro* CNS models to adequately recapitulate a breadth of human-specific pathologies is of utmost importance.

The majority of *in vivo* CNS disease study involve rodent models, allowing the observation of cellular phenotypes in a living system where cells are in their natural multicellular, 3D environment (Dawson, Golde, & Lagier-Tourenne, 2018; McGonigle, 2014). However, significant differences in the behavior and genetics of rodent models severely impacts the generalisability of these systems to human CNS diseases (McGraw, Ward, & Samaco, 2017). Contrastingly, most *in vitro* studies are conducted on 2D surfaces with monocultured cell lines, ignoring the 3D multicellular nature of the CNS (Birgersdotter, Sandberg, & Ernberg, 2005). This 2D monolayer culture causes significantly altered cell morphology, function, differentiation and gene expression compared to the *in vivo* environment, diminishing the physiological relevance of these cultures (Edmondson, Broglie, Adcock, & Yang, 2014; Sun, Jackson, Haycock, & MacNeil, 2006). The validity of these commonly used models for human CNS disorders is concerningly low (Liu, Xie, Meng, & Kang, 2019; Pound & Ritskes-Hoitinga, 2018) and as such, there is a need for further improvements to current *in vitro* culture systems.

The ability to create more relevant and increasingly complex *in vitro* models of the CNS environment has been revolutionized by the discovery of induced pluripotent stem cells (iPSCs) (Takahashi & Yamanaka, 2006). Cells differentiated from iPSCs provide a scalable solution for the production and culture of tissue-specific cell types which closely mimic their human *in vivo* counterparts and allow tighter temporal control of any experimental conditions (Heikkila et al., 2009; Johnson, Weick, Pearce, & Zhang, 2007; Zhang et al., 2017). iPSC-derived cerebral organoids exhibit extensive developmental and structural aspects of the CNS (Lancaster & Knoblich, 2014), however, this model system is plagued with low reliability and lack of control over cell-type ratios, diminishing its value for the high-throughput screening often required during pre-clinical therapeutic development (Di Lullo & Kriegstein, 2017; Quadrato et al., 2017).

In addition to the benefits provided by iPSC technology, hydrogel-based 3D cell culture represents a further biotechnological advancement in the development of biomimetic, replicable and scalable *in vitro* CNS models. Others have successfully cultured neural cells in a variety of naturally derived hydrogels consisting of one or more naturally occurring proteins, such as collagen (Egawa, Kato, Hiraoka, Nakaji-Hirabayashi, & Iwata, 2011) or Matrigel (a soluble mouse tumor basement membrane extract) (Choi et al., 2014), and synthetic poly(ethylene glycol) (PEG) hydrogels customised with cellular adhesion domains (Li, Zheng, Wang, Becker, & Leipzig, 2018). Natural hydrogels are often highly biocompatible in *in vitro* cell culture, however they suffer from batch-to-batch variability and often have a constrained relationship between their functional groups and biomechanical properties (Aisenbrey & Murphy, 2020). Additionally, natural hydrogels often have unfavorable rheological characteristics or gelation conditions for widescale bioprinting, hence are limited in their use (Mancha Sanchez et al., 2020).

In contrast, utilising synthetic PEG-based hydrogels allows for highly tunable biochemical and biomechanical properties, to accurately replicate the extracellular environment being studied. This tunability takes the form of protease-sensitive peptide sequences for biodegradation and cellular migration, covalent coupling of bioactive cellular adhesion sequences and changes in stiffness and porosity through variations in polymer chain length and concentration (Zhu, 2010). Additionally, efficient crosslinking within PEG-based hydrogels can be achieved through chemical/physical means without causing reductions in cellular viability (Li et al., 2018). These hydrogels are also highly compatible with high-throughput drop-on-demand printing (Utama et al., 2020).

Relevant *in vitro* CNS models should support viable and controlled growth of neurons, glia and vascular cells, exhibit relevant cell-type specific morphology and facilitate appropriate physiological responses to biological or pharmacological stimuli (Hopkins, DeSimone, Chwalek, & Kaplan, 2015). Challenges still remain for producing *in vitro* models with highly tunable scaffolds which mimic the extracellular matrix (ECM), support heterogenous tissue architecture and can be produced in a replicable, scalable and high-throughput format. Therefore, we aim to provide a customisable high-throughput 3D bioprinting platform that can successfully model essential aspects of the CNS environment *in vitro* using a defined and tunable physiologically- and mechanically-relevant matrix.

## 2. Materials and Methods

### 2.1 Cell culture and derivations

iPSC-derived astrocytes, BMECs and neurons followed methods previously published by (Bardy et al., 2015; Neal et al., 2019; Tcw et al., 2017). Specific details on cell culture and derivations can be found in the supplementary information.

### 2.2 Immunofluorescence

For 2D samples, cells were plated on Matrigel (0.08 mg/mL) coated glass chambers. 24 h post plating, cell media was aspirated and cells were washed 3 x with PBS. Cells were fixed with either 100 % ice-cold MeOH or 4 % paraformaldehyde in PBS (10 min, 22 °C). For MeOH protocol (used for 2D endothelial samples), samples were then rehydrated for 20 mins in PBS (10 min, 22 °C). For PFA, samples were permeabilized with 0.1% Triton-X-100 (10 min, 22 °C). Blocking was done in 5 % (v/v) FBS in PBS (blocking solution; 1 h, 22 °C). Primary antibodies were diluted in blocking solution (**Table S1**) and incubated overnight (4 °C). The following day, cells were washed 3 x with PBS and incubated with the appropriate Alexa 488/594-conjugated secondary antibody at 1:200 in blocking solution (1 h, 22 °C) (**Table S2**). Cells were washed 3 x in PBS and mounted on a glass coverslip in DAPI antifade solution (Sigma-Aldrich). Confocal microscopy was performed using the LSM800 (Zeiss, ZEN Blue software) and images processed using FIJI image analysis software.

For 3D samples, cells were fixed with 4 % paraformaldehyde in PBS (20 min, 22 °C), washed 3 x in PBS (10 min, 22 °C), then permeabilized with 0.1 % Triton-X-100 (20 min, 22 °C). Samples were then blocked with blocking solution (60 min, 22 °C) and primary antibodies were added in blocking solution (24 h, 4 °C). Samples were washed 3 x (5 min (22 °C), then 1 h (22 °C) then 24 h (4 °C)) with 0.1 % v/v Tween-20 in PBS (PBST) under gentle rocking. Samples were then incubated with secondary antibodies in blocking solution (24 h, 4 °C). Samples were washed 3 x in PBST as described above then incubated with 20 μM Hoechst (10 min), washed 1 x with PBS (22 °C) and then stored in PBS at 4 °C until imaged.

### 2.3 Matrix synthesis and 3D bioprinting

3D bioprinting was performed using the RASTRUM bioprinter from Inventia Life Sciences as described previously (Jung et al., 2022; Utama et al., 2021). During priming and printing of the inert base layer, cells were dissociated, centrifuged and resuspended in 200 μL of activator solution (Inventia Life Sciences) containing 3 mg/mL collagen IV at 10 x 10^6^ cells/mL for iBMECS and NPCs and 5 x 10^6^ cells/mL for astrocytes. Cell-laden matrices were printed in 96-well plates.

### 2.4 Viability/morphology analysis

Primary human astrocytes were cultured on Matrigel-coated (0.08 mg/well) 6-well plates in astrocyte growth medium (Lonza). On the day of plating, cells were dissociated using accutase (37 °C, 5 min), centrifuged (300 g, 5 min) and resuspended in activator (containing collagen IV (Sigma-Aldrich), Laminin 521 (BioLamina), both or neither) at 5 x 10^6^ cells/mL. 2 mL of the bioink (containing RGD, YIGSR, both or neither) was added into the center of a 96-well plate to form a small droplet on the bottom of each well. An equal volume of cell-containing activator (2 mL) was added directly into the droplet and allowed to gelate (22 °C, 2 min) before adding media and left to incubate (37 °C, 5% CO_2_).

To assess viability and morphology 24 h after plating the LIVE/DEAD viability/cytotoxicity kit (Invitrogen) was used. 3D cultures were washed 1 x with PBS and incubated in 2 μM calcein AM, 4 μM ethidium homodimer-1 and 20 μM Hoechst 33342 (Sigma-Aldrich) (37 °C, 30 min). Wells were washed 3 x with PBS and fluorescent microscopy was achieved using the Cytation 3 Cell Imaging Multi-Mode Reader (BioTek, Gen5 v2 software), with a combination of 375 nm, 480 nm and 560 nm excitation filters. Images were taken as 50 z-sections spaced at 2 μm intervals on a 10 x air objective, each condition was plated in duplicate and a minimum of 80 cells were imaged. Images were processed as maximum intensity projections in FIJI. For analysis, images were blinded and cell area and maximum process length were measured by manual tracing.

### 2.5 IL-6 ELISA

iPSC derived astrocytes were plated on Matrigel-coated (0.08 mg/well) 96-well plates or 3D-printed to achieve a density of 15,000 cells/well in both conditions. 24 h after plating, media was aspirated and cells were exposed to 10/50 μg/mL Lipopolysaccharide (LPS; *E. coli* strain 0111:B4; Sigma-Aldrich) (MilliQ H_2_O vehicle) and 50 μg/mL TLR4 blocking antibody (14-9917-82; Invitrogen) in astrocyte medium for 24 h. Media was removed, centrifuged (10,000 *g*, 1 min) and the supernatant stored at −80 °C. IL-6 ELISA was performed according to manufacturer’s protocols (R&D Systems product number DY206), with slight amendments including: capture antibody used at 2 μg/mL, detection antibody used at 50 ng/mL and standard range used from 0.586-600 pg/mL. Absorbance at 450 nm was recorded using a BMG POLARstar Omega and analysed using a sigmoidal dose-response (variable slope) model, with unknowns interpolated in GraphPad Prism 9.

### 2.6 Calcium Imaging

After 4 weeks of differentiation, iPSC-derived neurons were washed once with BrainPhys basal media and incubated with Fluo-4 calcium dye (1:1000), probenecid (1:250) and PowerLoad concentrate (1:100) (Fluo-4 Calcium Imaging Kit, Molecular Probes) in BrainPhys basal media for 30 min (37 °C, 5 % CO_2_) then 15 min (RT). Fluo-4 containing media was removed, cells washed 1 x and left in BrainPhys basal media. The cells were then immediately imaged for 10 min at a rate of 1 frame every 0.6 sec using the LSM800 under a 10 x air objective and live cell conditions (37 °C, 5 % CO_2_). The analysis of calcium images was done in MATLAB (R2021b) using the ‘EZCalcium’ toolbox developed and validated by (Cantu et al., 2020), which automates the region of interest selection and generates calcium traces for each neuron body identified. These calcium traces were then quantified using the ‘PeakCaller’ MATLAB script developed by (Artimovich, Jackson, Kilander, Lin, & Nestor, 2017).

## 3. Results

### 3.1 Matrix optimisation

To identify a suitable CNS-mimetic matrix composition, we utilised PEG-based matrices with stiffnesses similar to that of the human cortex (1.1 and 1.5 kPa) (Weickenmeier et al., 2016) and screened a small selection of functional adhesion peptide motifs and proteins. Although ultimately we are seeking to develop a matrix suitable for bioprinting, bioprinting can impact the viability of cells due to the shear stress cells are exposed to (Shi et al., 2018). Given this, to assess the effect of matrix composition on astrocyte viability and morphology in the absence of bioprinting-related stressors, we examined the impact of manually encapsulating (pipetting) primary human foetal astrocytes in a range of PEG-based matrices containing various combinations of adhesion molecules/peptides. We incorporated adhesion peptide motifs found within fibronectin (RGD sequence) and laminin (YIGSR sequence) as well as the full-length collagen IV and laminin proteins at concentrations consistent with previous literature (Antoine, Vlachos, & Rylander, 2014; Pereira, Barrias, Bartolo, & Granja, 2018; Swindle-Reilly et al., 2012).

In the 1.1 kPa matrix, we found all of the functionalised matrix conditions to significantly increase the viability of primary human astrocytes compared to the unfunctionalised blank matrix (**Figure 1 A**). We found similarly significant results within the 1.5 kPa matrix, with the exception of YIGSR where it showed no significant increase over the blank matrix (**Figure 1 D**). We then examined the extent to which astrocytes adhered to various matrix conditions by quantifying the average cell size and the maximum distance in which each cell was extending into the matrix. In contrast to cell viability, we found the different matrix compositions in both the 1.1 kPa (**Figure 1 B and C**) and 1.5 kPa (**Figure 1 D and E**) matrices to have a much more varied effect on astrocyte adherence. Incorporation of RGD and collagen IV both alone and in combination caused significant increases to cell area compared to the blank matrix regardless of stiffness (**Figure 1 B and E**). Unique to the 1.5 kPa matrix, the addition of laminin to the combination of RGD + collagen IV as well as the combination of RGD + collagen IV + YIGSR resulted in no significant increase in cell extension compared to the blank matrix (**Figure 1 E and F**).

**Figure 1.**
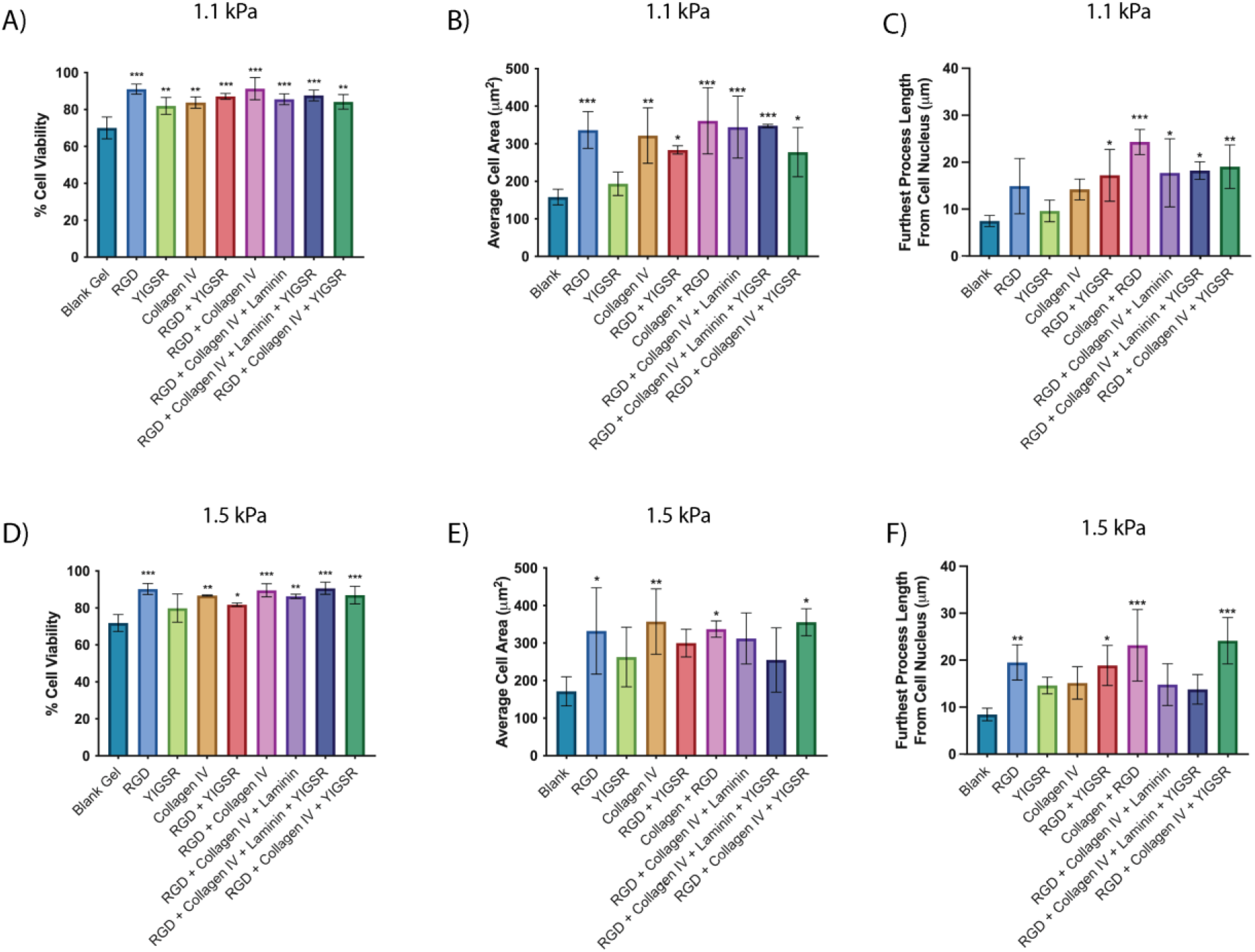
Figure shows the viability, average cell area and average longest cellular process of primary human fetal astrocytes one day post-encapsulation in a 1.1 kPa PEG-based matrix (A-C) and 1.5 kPa PEG-based matrix (D-F) containing various adhesion peptides and proteins (RGD, YIGSR, collagen IV and laminin). Data shows the mean ± SD of n ≥ 3 independent experiments. A one-way ANOVA with Dunnet’s multiple comparison test was used to determine whether there were statistically significant differences between each peptide/protein containing matrix and the blank matrix. (* p < 0.05, ** p < 0.01, *** p < 0.001).

Despite the matrix optimisation being based upon primary human astrocyte biocompatibility, the lead matrix will ultimately facilitate a 3D culture system that aims to support multiple CNS cell types. RGD and collagen IV-functionalised matrices provided the highest level of biocompatibility for the primary astrocytes, however despite showing no benefit to astrocyte size or process extension, YIGSR benefits the growth of other CNS cell types such as neurons and brain microvascular endothelial cells (BMECs) (Grant et al., 1989; Jain & Roy, 2020). Selecting a lead matrix composition which incorporates the widest variety of motifs available, namely RGD, YIGSR and full-length collagen IV provides a better opportunity for other CNS cell types to be compatible with the matrix. As such, given that the RGD + collagen IV and RGD + collagen IV + YIGSR conditions performed equally as well in the 1.5 kPa matrix and exhibited slightly higher cell adhesion properties compared to the 1.1 kPa matrix, we chose the 1.5 kPa RGD + collagen IV + YIGSR composition as our lead matrix to use for further applications. A simple schematic of this matrix composition is found in

### 3.2 Bioprinted iPSC-derived astrocytes remain viable and respond differently to inflammatory stimuli

Utilising iPSC-derived CNS cells in conjunction with 3D matrices and bioprinting technologies provides an opportunity for the production tissue-specific cell types within increasingly complex *in vitro* models of the CNS. We report the successful differentiation of iPSC-derived astrocytes following a previously published protocol (Tcw et al., 2017) which express the astrocytic markers S100ß and GFAP (**Figure 3 A**). We then 3D-printed these iPSC-derived astrocytes with the 1.5 kPa RGD + collagen IV + YIGSR matrix using a commercial 3D bioprinter. iPSC-derived astrocytes were printed with good viability (82.2 % ± 10.1 SD) and exhibited extensive hypertrophy of cell bodies and stretched processes characteristic of astrocyte morphology one day post-printing (**Figure 3 B and C**). It’s worth noting that 3D-printed iPSC-derived astrocytes exhibited GFAP staining at similar levels to background compared to the intense staining observed in 2D cultured astrocytes (**Figure 3 A and C**).

**Figure 2.**
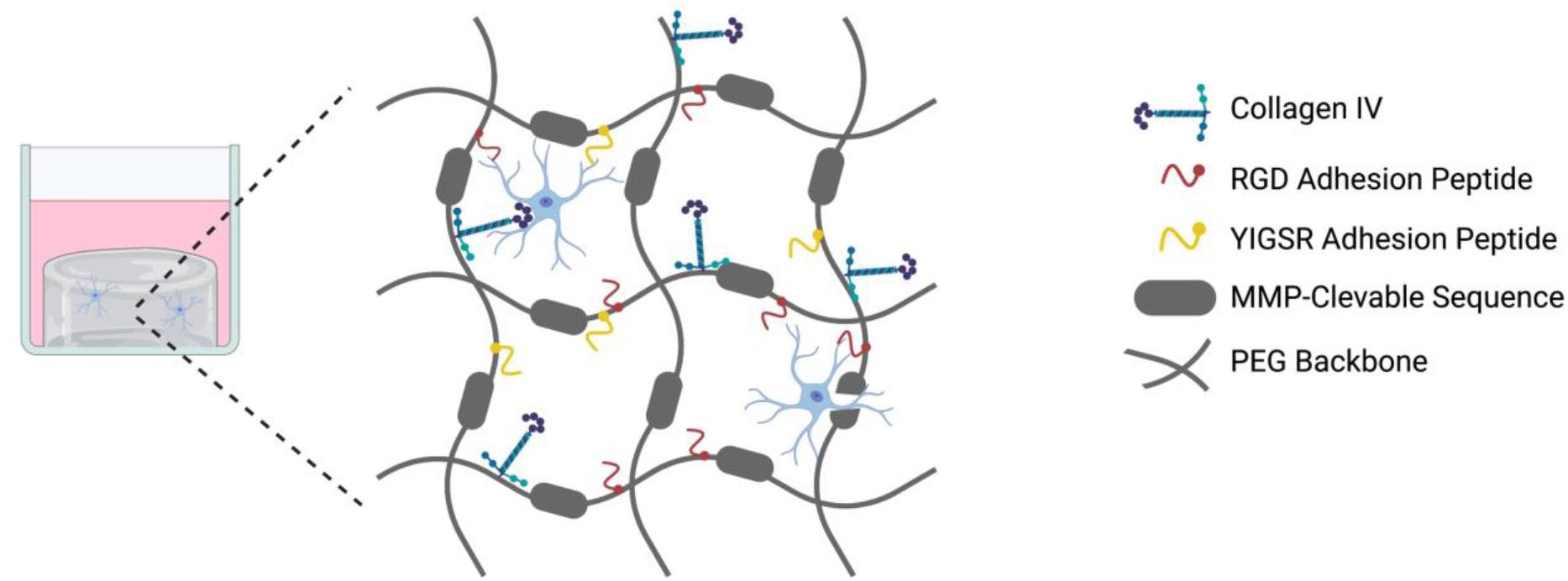
Simplified schematic of the matrix composition showing the functionalisation of the matrix backbone with collagen IV, Arg-Gly-Asp (RGD) peptide sequence and Tyr-Ile-Gly-Ser-Arg (YIGSR) peptide sequence. Created with BioRender.com.

**Figure 3.**
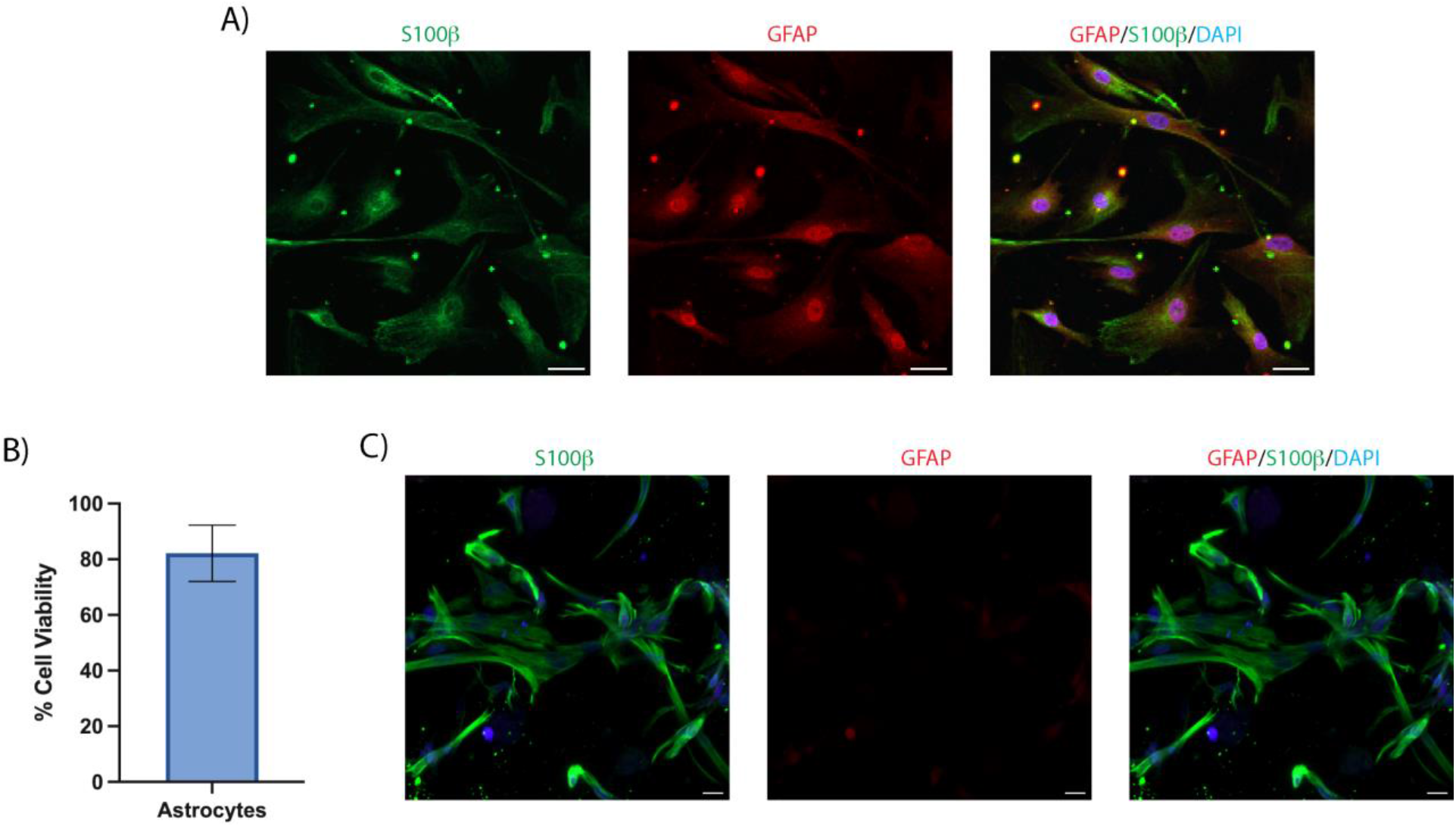
A) Immunocytochemistry of iPSC-derived astrocytes in 2D culture and stained for the astrocyte markers S100ß (green), GFAP (red) and nuclei counterstained with DAPI (blue). B) Viability of iPSC-derived astrocytes one day post-bioprinting, data shows the mean ± SD of n = 3 independent experiments. C) Immunocytochemistry of iPSC-derived astrocytes bioprinted in the 3D matrix and shows a maximum intensity projection of an 80 μm z-stack. Scale bars = 20 μm.

Successful 3D *ïn vitro* models of the CNS should be constructed from an immunologically inert matrix scaffold and facilitate neuroinflammatory responses, of which astrocytes play a significant role (Giovannoni & Quintana, 2020). We aimed to characterise the biological response to an inflammatory stimulus and pharmacological intervention in 2D cultured and 3D bioprinted iPSC-derived astrocytes. We found no significant difference in the basal IL-6 secretion between 2D and 3D bioprinted astrocytes (**Figure 4**). LPS is a bacterial-derived polysaccharide which induces proinflammatory cytokine release through the activation of the Toll-like receptor 4 (TLR4) receptor (Gorina, Font-Nieves, Marquez-Kisinousky, Santalucia, & Planas, 2011). Upon both 10 and 50 μg/mL LPS stimulation, astrocytes in 2D significantly increased their IL-6 secretion from basal. Compared to astrocytes cultured in 2D, 3D bioprinted astrocytes showed significantly less IL-6 production in response to both 10 and 50 μg/mL LPS (**Figure 4**). A TLR4 blocking antibody reduced the IL-6 response from 2D astrocytes in response to 10 and 50 μg/mL LPS, albeit only statistically significant for 10 μg/mL LPS. Conversely, 3D bioprinted astrocytes showed no change in LPS-stimulated IL-6 secretion in response to the TLR4 blocking antibody (**Figure 4**).

**Figure 4.**
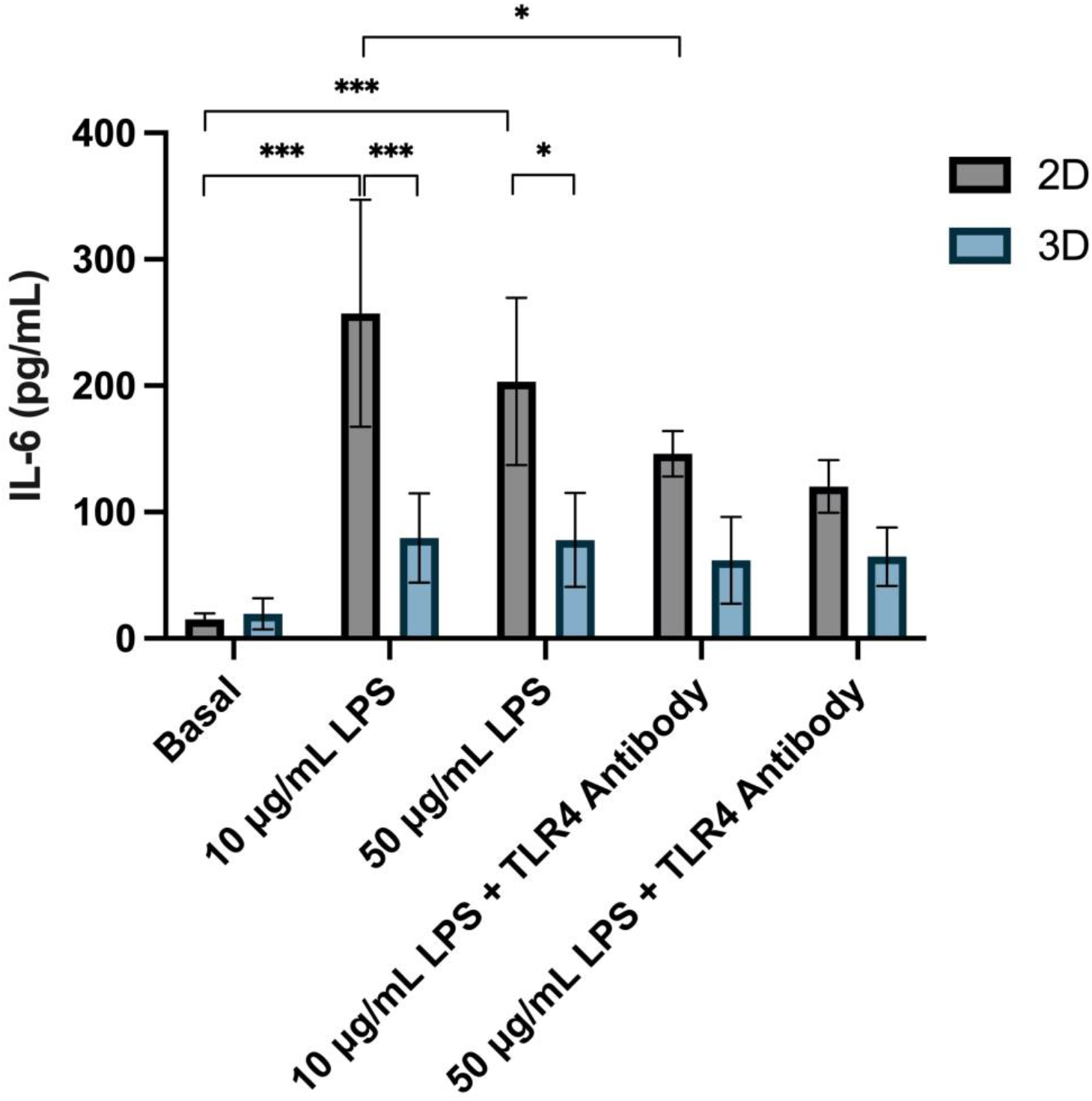
Concentration of IL-6 secreted from 2D and 3D bioprinted iPSC-derived astrocytes basally and after exposure to 10 or 50 μg/mL LPS and 50 μg/mL TLR4 blocking antibody for 24 h. Data shows the mean ± SD of n ≥ 3 independent experiments. A two-way ANOVA with Tukey’s multiple comparison test was used to determine significant differences between conditions, relevant comparisons where a significant difference was found are shown (* p < 0.05, *** p < 0.001).

### 3.3 Bioprinted iBMECs remain viable and undergo vasculogenesis

iPSC-derived BMECs (iBMECs) were validated by immunofluorescence for expression of common endothelial markers (**Figure S1 A**). After derivation, iBMECs were then bioprinted or seeded onto 2D fibronectin/collagen IV-coated wells. One day after printing in 2D, iBMEC viability was measured using a Cell Titre Blue assay (CTB). When cultured in the suggested post-derivation media from Neal et al. (2019) (hESFM with 0.5 % B27 supplement, herein referred to as endothelial medium (EM)), 2D bioprinted iBMECs showed similar viability to non-printed iBMECs plated manually with a pipette (**Figure 5 A**). However, culturing in ScienCell Astrocyte Medium (AM) increased the viability of 2D bioprinted iBMECs compared to EM at day 1. Additionally, by applying 10 μM of the ROCK inhibitor Y-27632 (ROCKi), cell viability of 2D bioprinted cells was increased in both EM and AM conditions, with AM + ROCKi supporting the highest viability cultures (**Figure 5 A**). After 7 days, good viability was evidenced in all AM-containing conditions, which reached 100 % confluency, while EM conditions showed markedly lower viability (**Figure 5 B**).

**Figure 5.**
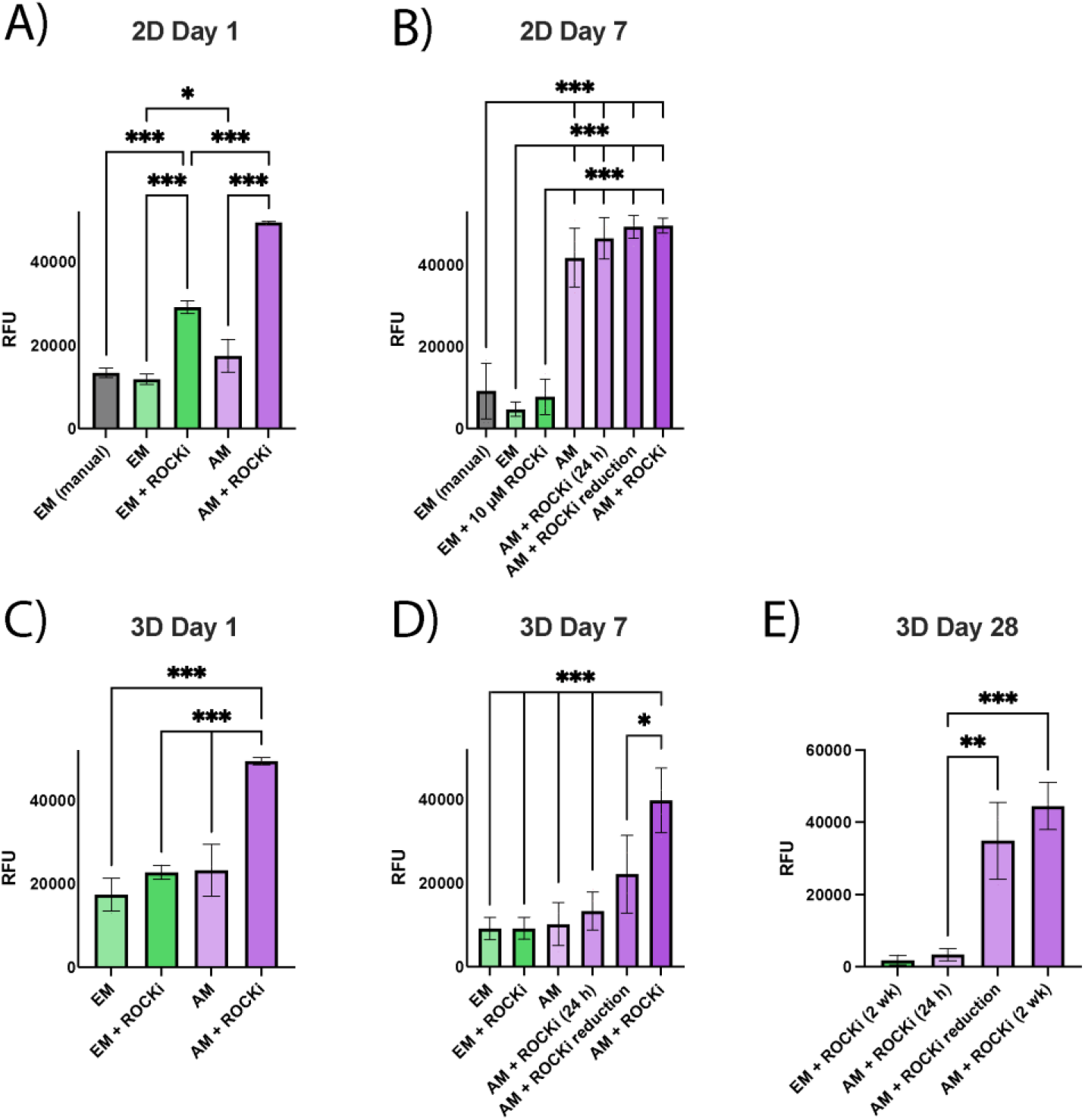
Viability of human iPSC-derived brain microvascular endothelial cells (iBMECs) bioprinted in 2D or 3D matrices under different media conditions. Comparison of raw fluorescence units (RFU) after 4 h incubation with Cell Titre Blue in 2D cultures at day 1 (A) and day 7 (B), and in 3D cultures at day 1 (C), day 7 (D) and day 28 (E). Graphs are presented as mean ± SD of n ≥ 3 independent experiments. A-E are compared using a one-way ANOVA with Tukey’s multiple comparisons test. (* p < 0.05, ** p < 0.01, *** p < 0.001).

iBMECs printed in 3D PEG matrices in comparison to 2D, displayed cell viability that was more media-dependant. At day 1, CTB results demonstrated that AM + ROCKi provided significantly higher cell viability than EM and AM without ROCKi (**Figure 5 C**). iBMECs cultured in AM + ROCKi displayed a sufficient percentage of viable cells (67.71 % ± 11.28 SD). iBMECs in EM + ROCKi showed similar viability (56.75 % ± 14.67 SD) but contained a high proportion of atypical nuclear staining that was delocalised from either calcein AM (live) or EthD (dead) staining (**Figure S1 B - D**). This mirrors CTB results that indicate EM provides sub-optimal conditions for iBMEC survival in this system. In the two conditions that did not contain ROCKi, calcein AM staining was rarely associated with a typical live cell morphology or appropriate nuclear staining (**Figure S1 C**). Because of this, live cell counts were impractical, but this atypical live/dead finding corroborates CTB results suggesting poor viability in these conditions.

At day 7, 3D bioprinted iBMECs cultured in AM + ROCKi retained high viability measured by CTB, while those cultured in EM (with or without ROCKi), and AM without ROCKi retained their comparably low viabilities (**Figure 5 D**). Typically, ROCKi is removed after 24 h in culture (AM + 24 h ROCKi), however iBMEC viability significantly declined after the removal of ROCKi 24h post-bioprinting, compared to culturing in AM with daily ROCKi replacements, which had significantly higher viability than all other conditions. Unfortunately, ROCKi application may have deleterious effects on endothelial cell pathways (Cao et al., 2017). Therefore, a reduction scheme was devised to limit exposure, whereby ROCKi concentration was reduced by 2 μM per day starting on day 1 (AM + ROCKi reduction). At day 7, AM + ROCKi reduction was not significantly different from AM + ROCKi (24 h). By day 28, however, iBMECs cultured in AM with 2 weeks ROCKi, and the ROCKi reduction scheme show comparably high viability, compared to the significantly poorer viability in EM + ROCKi and AM + ROCKi (24 h) conditions (**Figure 5 E**). This is corroborated by live cell counts, showing similar live cell percentages for AM + ROCKi (66.58 % ± 23.67 SD) and AM + ROCKi reduction (75.94 % ± 3.68 SD) (**Figure S1 E**).

At day 28, these cultures expressed markers and displayed cell morphology similar to *in vivo* vascular formations, progressing from small, rounded, singularized cells at day 1, to elongated multicellular structures that mimic *in vivo* vascular networks at day 28 (**Figure 6 A**). These iBMEC networks were stained for the adherens junctions protein VE cadherin, the adhesion protein, PECAM-1, the tight junction protein, occludin, the extracellular matrix protein, laminin α4, and the functional glucose transporter, GLUT1, demonstrating that endothelial cell identity is maintained (**Figure 6 B**). The structural complexity of the vascular networks is highlighted by staining of F-actin (**Figure 6 C**).

**Figure 6.**
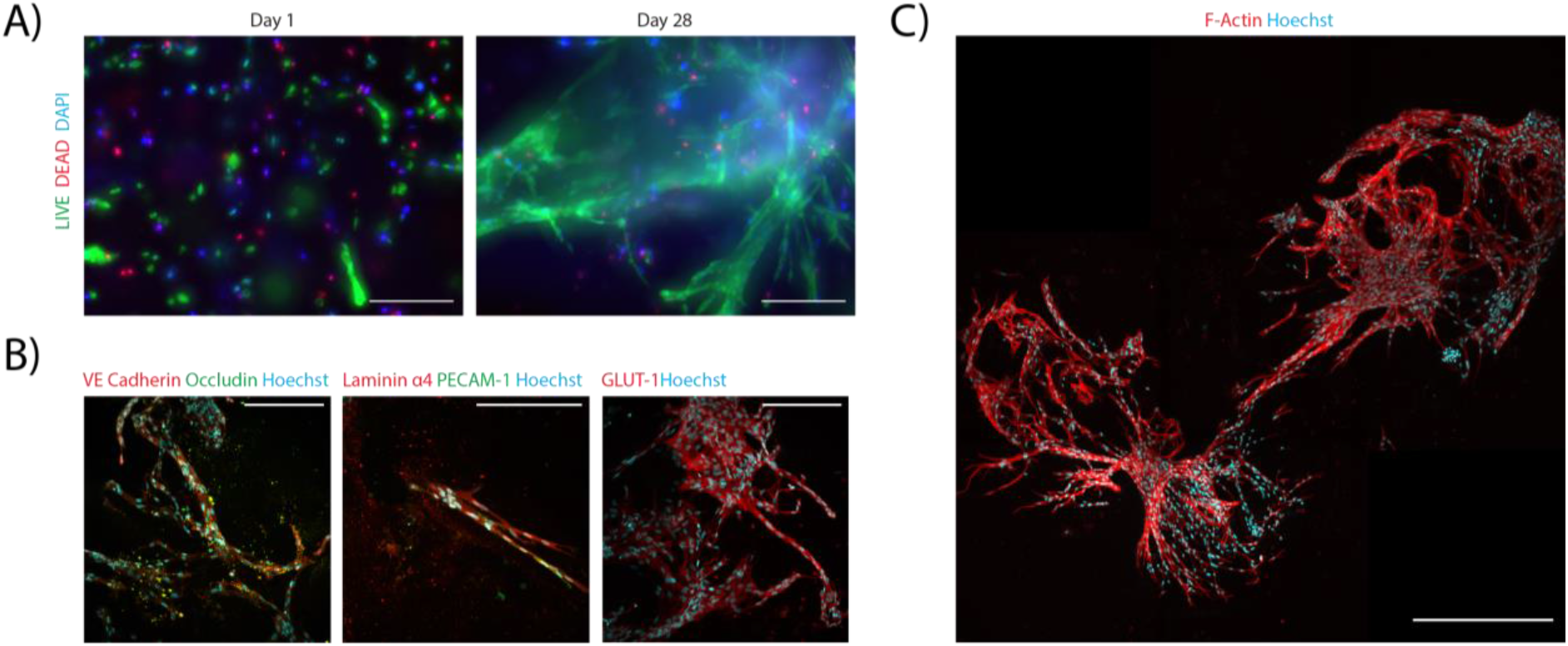
Immunofluorescence images of 3D bioprinted iBMECs on day 28. A) Maximum intensity projections of iBMECs at Day 1 and Day 28, stained with calcein AM (live) and EthD (dead). Images taken using a 10 × objective. Scale bar = 200 μm B) Maximum intensity projections of the expression of VE cadherin, occludin, laminin α4, PECAM-1 and GLUT-1 by iBMECs. Images taken using a 25 × water objective. Scale bar = 200 μm C) Maximum intensity projection of F-actin staining of iBMECs shows the development of a complex network throughout the matrix. Image taken using a 40 × dry objective. Scale bar = 500 μm.

### 3.4 Bioprinted NPCs remain viable and enhances neuronal differentiation

Neurons are the most widely studied cell type within the brain and are an essential aspect within *in vitro* models of the CNS. NPCs were able to be successfully bioprinted within our lead matrix with good viability one-day post printing (82.2 % ± 9.2 SD) (**Figure 7 B**) and were differentiated into neurons after 4 weeks (**Figure 7 A**). Detection of the mature neuronal marker MAP2 was used to characterise the extent of neuronal differentiation between 2D and 3D culture conditions. We found differentiating neurons within 3D culture significantly increased in the percentage of MAP2 positive compared to 2D (**Figure 7 C**), however we observed no significant difference in the average neurite extension of MAP2 positive cells between 2D and 3D cultures (**Figure 7 D**).

**Figure 7.**
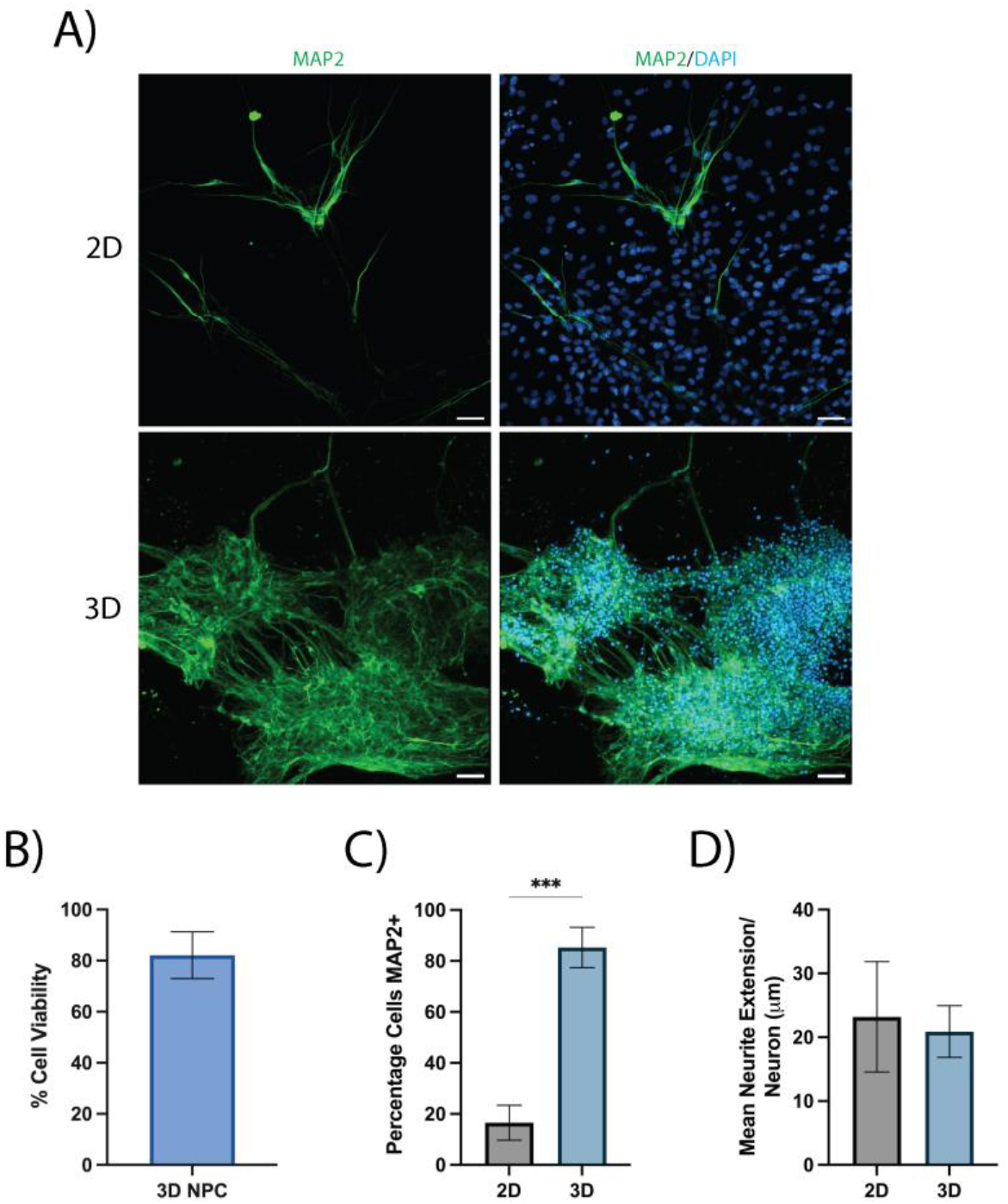
A) Immunofluorescence images of iPSC-derived neurons cultured in 2D and 3D bioprinted cultures after 4 weeks of differentiation. The cells were stained for the mature neuronal marker MAP2 (green) and all nuclei were counterstained with DAPI (blue). 3D culture images are represented as a maximum intensity projection of a 100 μm z-stack. Scale bars = 50 μm. B) Viability of NPCs one day post bioprinting. C) Percentage of iPSC-derived neurons positive for MAP2 and D) average neurite extension per neuron after differentiation for 4 weeks in either 2D or bioprinted 3D cultures. Data shows the mean ± SD of n = 3 independent experiments, an unpaired t-test was used to determine significant differences between 2D and 3D conditions. (*** p < 0.001).

To generate physiologically relevant *in vitro* models of the CNS, 3D matrices should facilitate and support spontaneous neuronal activity. We performed a quantitative and automated analysis of live-cell calcium imaging in both 2D and 3D bioprinted cultures of iPSC-derived neurons following two previously published protocols (Artimovich et al., 2017; Cantu et al., 2020). In comparison to 2D culture, we found iPSC-derived neurons cultured in 3D showed a significant increase in the number of spontaneous calcium spikes per neuron (**Figure 8 A**) and a significant decrease in the proportion of neurons producing spontaneous calcium spikes (**Figure 8 D**). There was no significant difference between 2D and 3D cultures in either the average signal height or the maximum signal height (**Figure 8 B and C**).

**Figure 8.**
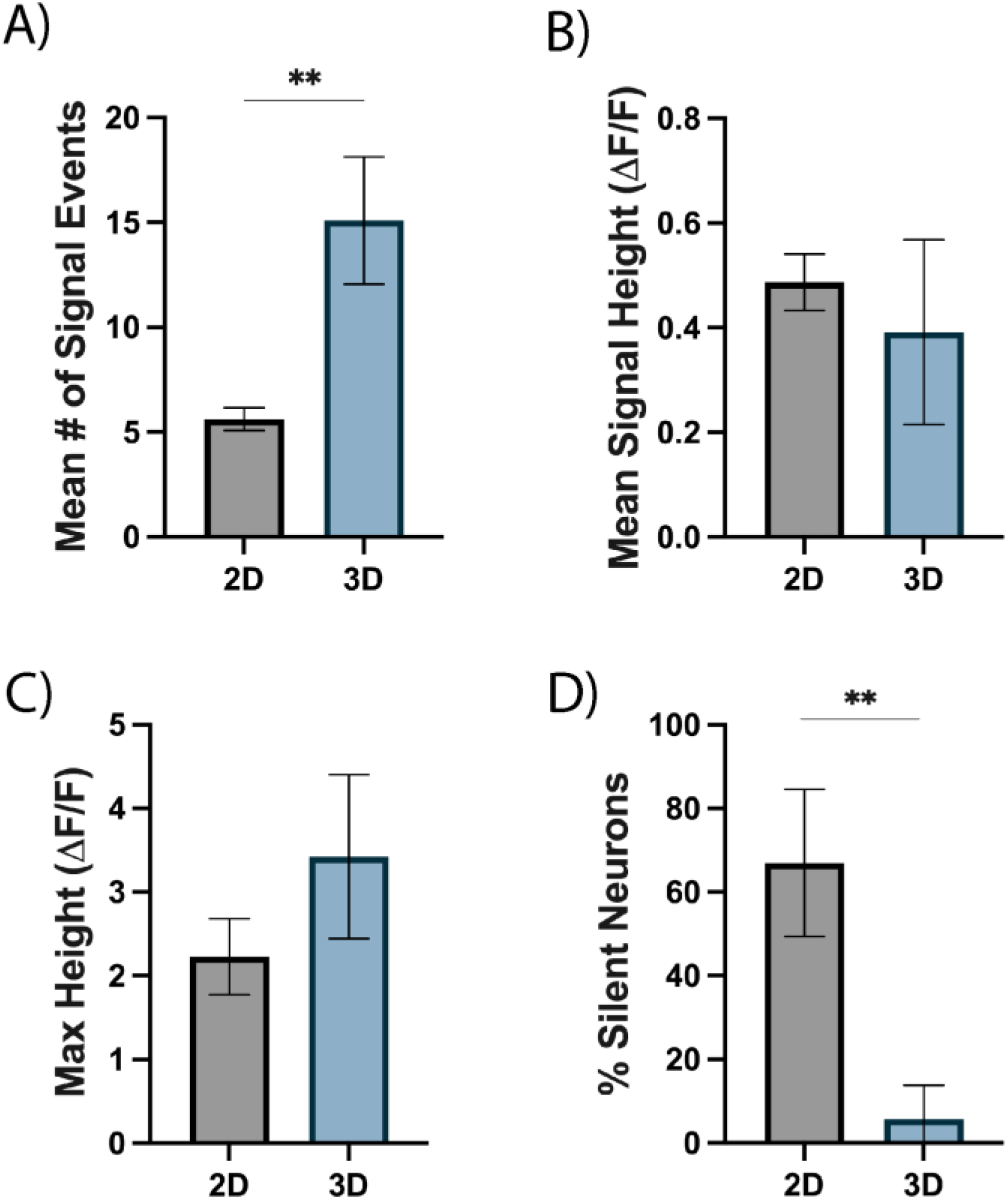
Quantification of spontaneous live-cell calcium imaging of iPSC-derived neurons cultured in 2D and bioprinted 3D cultures after 4 weeks of differentiation. The figure shows the quantification of A) mean number of spontaneous calcium transients per active neuron, B) mean signal height of each spontaneous calcium transient as measured by the change in fluorescence (ΔF/F), C) mean maximum height of calcium transients in each neuron as measured by ΔF/F and D) proportion of neurons which showed no spontaneous calcium transients (termed silent neurons) over the recording period (10 min). An unpaired t-test was used to determine significant differences between conditions. (** p < 0.01).

## 4. Discussion

Developing physiologically relevant *in vitro* models of the CNS is crucial in furthering our understanding of disease pathogenesis and facilitate the screening of potential therapeutics. The success of these models depends on the ability to support the growth and function of multiple physiologically relevant CNS cells. Here we report the optimisation of a novel CNS-mimetic 3D matrix and supports viable culture of 3D bioprinted iPSC-derived astrocytes, iBMECs, NPCs and enhanced neuronal differentiation and function.

During matrix optimisation, we showed that regardless of matrix stiffness, addition of RGD or collagen IV to our 3D PEG matrix provided benefits to the adhesion and process extension of primary human astrocytes. The benefit of RGD and collagen incorporation within 3D matrices is consistent with a number of previous studies culturing CNS cells (Mauri et al., 2018; Scott, Marquardt, & Willits, 2010). Despite concerns surrounding reductions in cell viability due to shear stressors during bioprinting (Shi et al., 2018), we were able to successfully bioprint iPSC-derived astrocytes, NPCs and iBMECs within our lead matrix with a good level of viability similar to what we found using manual gelation. Recent work using extrusion-based bioprinting of rodent astrocytes within composite gelatin-methacryloyl matrices and NPCs within collagen-based matrices have reported viability ranging from 75 - 90 %, similar to our results (de Melo, Cruz, Ribeiro, Mundim, & Porcionatto, 2021; Ouyang et al., 2020; Salaris et al., 2019; Sharma, Smits, De La Vega, Lee, & Willerth, 2020). Furthermore, we show that iBMECs bioprinted in our lead matrix, cultured in AM with tapered ROCK inhibition, remain viable up to 4 weeks. While ROCK inhibition is used *in vitro* for the differentiation, expansion and protection of brain endothelial cells (Joo et al., 2012; Niego et al., 2017), it has been suggested to affect VE cadherin junctions and actin/myosin dynamics (Cao et al., 2017). After 28 days, iBMECs cultured under the exposure-limiting ROCKi reduction scheme showed comparable viability, marker expression and morphology to iBMECs cultured with 2 weeks of ROCKi. Maintained expression of junctional proteins (PECAM-1, VE Cadherin and Occludin) and functional proteins (the ECM protein laminin α4, and the glucose transporter GLUT-1) after 28 days indicate continued endothelial-identity and function throughout longitudinal culture.

In comparison to iPSC-derived astrocytes cultured in 2D, we observed a reduction in the immunofluorescent intensity of GFAP staining in 3D bioprinted astrocytes. This was accompanied by a significantly lower production of IL-6 upon LPS stimulation. Previous literature culturing primary and stem-cell derived astrocytes have shown that matrix and ECM composition greatly effects the level of astrocytic activation, GFAP expression (a widely used marker for astrocyte activation) and response to various stimuli (Escartin, Guillemaud, & Carrillo-de Sauvage, 2019; Johnson et al., 2007). Similar to our findings, human stem cell derived-astrocytes cultured within collagen-based and hyaluronic acid matrices showed reductions in GFAP expression in comparison to 2D culture (Galarza, Crosby, Pak, & Peyton, 2020; Placone et al., 2015; Seidlits et al., 2019). It’s possible that the lower GFAP intensity and less exaggerated pro-inflammatory response to LPS observed within our 3D bioprinted iPSC-derived astrocytes is reflective of a lower level of astrocyte activation. TLR4 expression has shown to change between activated astrocytes and may contribute to the reduction of LPS-stimulated IL-6 secretion in 2D astrocytes and lack of response to the TLR4 antagonist (Wu et al., 2012).

Many current 3D endothelial models are based on manual seeding, or techniques which pre-define the initial cellular architecture (e.g. sacrificial networks or subtractive fabrication), ignoring the importance of morphogenic self-assembly for *in*-*vivo*-relevant vascular modelling (Brassard & Lutolf, 2019; Grebenyuk & Ranga, 2019). Extensive vasculogenic self-assembly displayed in our 3D bioprinted iBMECs is reminiscent of iBMEC cultures manually seeded in Matrigel, fibrin and GelMA matrices (Blanchard et al., 2020; Calderon et al., 2017), supporting the suitability of this platform for investigating iBMECs function and morphology.

To further validate our 3D bioprinted model for *in vitro* cell modeling of the CNS, we characterised the ability of our model to facilitate efficient neuronal differentiation and support neuronal function. The enhanced neuronal differentiation we observed in 3D bioprinted cultures supports previous literature using other natural and synthetic-based matrices (Brannvall et al., 2007; Pellett et al., 2015; Ranjan et al., 2020). The enhancement of neuronal function in 3D culture is in line with previous reports of primary rodent neurons within a number of 3D matrices and found the signalling patterns to be more reflective of *in vivo* neuronal activity (Bosi et al., 2015; Bourke et al., 2018). Taken together, 3D bioprinting NPCs within our matrix provided a highly biocompatible environment which enhances the generation and function of iPSC-derived neurons.

Taken together, the current study showcases the ability of our 3D bioprinted cultures to facilitate a range of biocompatibility and functionality of iPSC-derived CNS cells. Furthermore, the tunability and scalability of this 3D bioprinted system makes it an attractive platform to further develop *in vitro* models of CNS disorders. A key area for further development of our platform involves the co-culture of multiple cell types. Co-culturing neurons with various glial cells elicits cellular functions more reflective of the *in vivo* environment (Goshi, Morgan, Lein, & Seker, 2020). As such, co-culturing iPSC-derived neurons, astrocytes, BMECs as well as other CNS cells such as oligodendrocytes and microglia within the scalable 3D platform presented within the current study is the logical next step for developing more complex *in vitro* 3D models. An important consideration for these more complex multi-cellular models for use in pre-clinical drug screening is ensuring cell-type ratios are consistent and reflective of the *in vivo* environment being modeled.

## Declarations

### • Ethics approval and consent to participate

Approval for the use of iPSCs was gained from the University of Sydney Institutional Biosafety Committee.

### • Data availability statement

The datasets used during the current study are available from the corresponding authors on reasonable request.

### • Authors’ contributions

M.A.S and S.D.L contributed to the design of the work, acquisition of data, analysis of data, interpretation of data and drafted the manuscript. A.V and M.E contributed to the design and revised the work. E.L.W conceived the work, contributed to the design of the work, contributed to the cell line acquisition and derivations, the interpretation of data and revised the work. M.K contributed to the conception of the work and revised the work. All authors have read and approved the submitted version.

## • Acknowledgements

The authors acknowledge the technical and scientific assistance of Sydney Microscopy & Microanalysis, the University of Sydney node of Microscopy Australia. We also would like to acknowledge Cedars-Sinai Medical Center’s David and Janet Polak Foundation Stem Cell Core Laboratory providing the iPSC lines used within the study.

## Supplementary Information

**Table S1.** Primary antibodies used in immunofluorescence images

| Target | Host | Vendor | Product Number | Dilution |
| --- | --- | --- | --- | --- |
| GFAP | rabbit | Abcam | ab7260 | 1:500 |
| S100β | mouse | Sigma-Aldrich | S2532 | 1:1000 |
| Nestin | mouse | Stem Cell | 60091 | 1:2000 |
| Pax6 | rabbit | Abcam | ab5790 | 1:50 |
| MAP2 | mouse | Invitrogen | MA5-12823 | 1:100 |
| Ve<br>Cadherin | rabbit | Abcam | ab33168 | 1:400 |
| Occludin | mouse | ThermoFisher | OC-3F10 | 1:100 |
| PECAM-1 | mouse | Abcam | ab9498 | 1:1000 |
| Laminin<br>α4 | rabbit | Sigma-Aldrich | SAB4501719 | 1:100 |
| GLUT-1 | rabbit | Abcam | ab14309 | 1:250 |

**Table S2.** Secondary antibodies used in immunofluorescence images

| Species<br>Reactivity | Host | Conjugate | Vendor | Product<br>Number | Dilution |
| --- | --- | --- | --- | --- | --- |
| Mouse | Donkey | Alexa Fluor 488 | ThermoFisher | A-21202 | 1:200 |
| Rabbit | Donkey | Alexa Fluor 594 | ThermoFisher | A-21207 | 1:200 |

**Figure S1.**
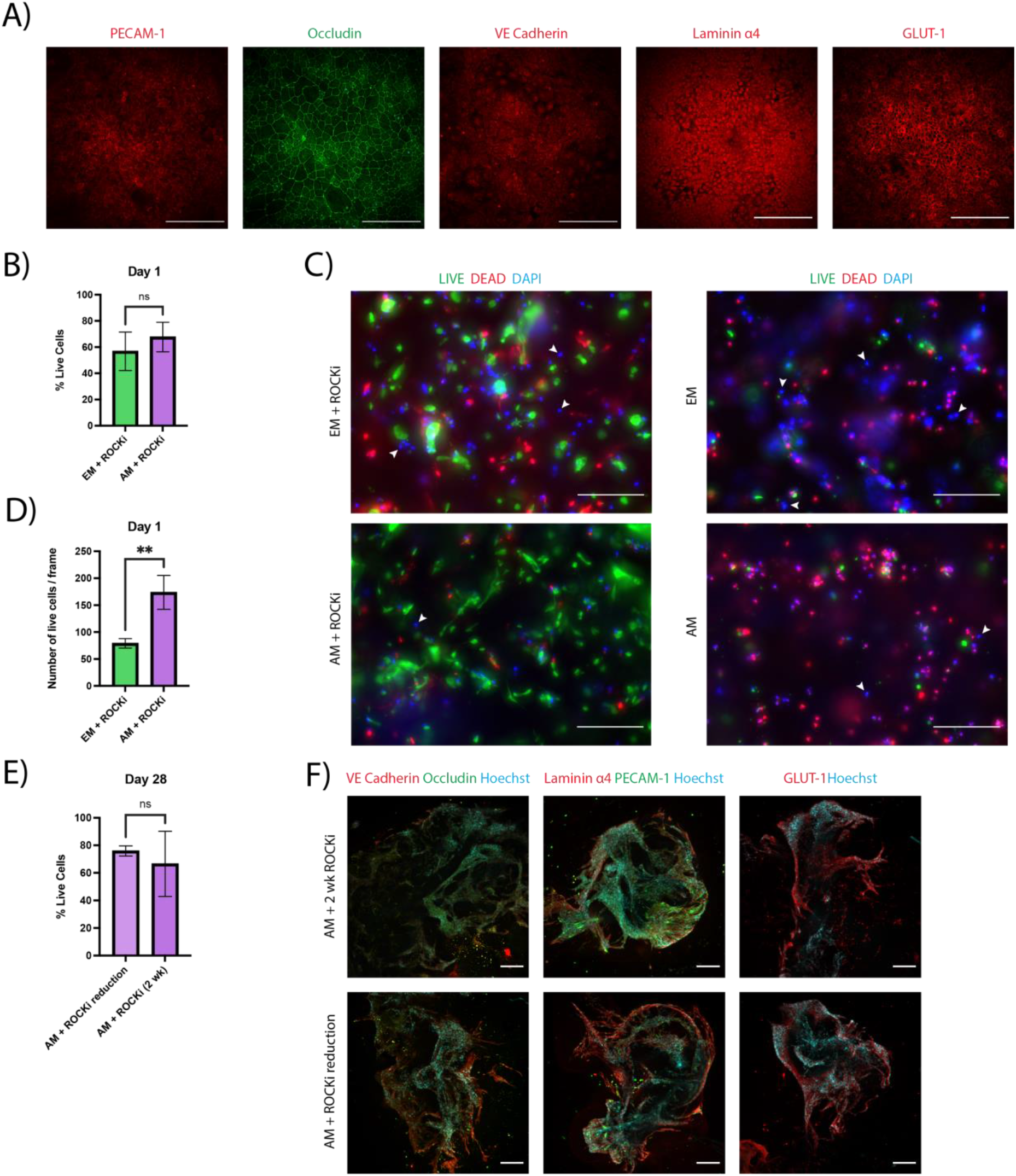
A) 2D iBMECs express VE cadherin, occludin, laminin α4, PECAM-1 and GLUT-1. Images taken using a 25 × water objective. Scale bar = 200 μm. B) Percentage of live cells at Day 1 identified by Live/Dead staining. C) Maximum intensity projections of iBMECs at Day 1 under different media conditions, stained with calcein AM and EthD. White arrows indicate examples of atypical nuclear staining. Images taken using a 10× objective. Scale bar = 200 μm. D) Number of countable live cells at Day 1 identified by Live/Dead staining. E) Percentage of live cells at Day 1 identified by Live/Dead staining. F) Maximum intensity projections of 3D iBMECs at Day 28 under different media conditions showing expression of VE cadherin, occludin, laminin α4, PECAM-1 and GLUT-1. Images taken using a 10× dry objective. Scale bar = 200 μm. Graphs are presented as mean ± SD. B, D and E are compared using an unpaired t-test. (* p < 0.05, ** p < 0.01, *** p < 0.001).

### iPSC Line and Culture

Human control iPSCs were obtained from the Cedars-Sinai (Los Angeles, USA) cell repository. iPSCs were generated from dermal fibroblasts obtained from skin punch biopsies and reprogrammed using a non-integrating episomal plasmid and showed normal karyotyping. iPSCs were cultured in feeder free conditions on Matrigel-coated (0.08 mg/well) 6-well tissue culture plates in mTesR1 media (StemCell Technologies) supplemented with mTeSR1 5× Supplement (StemCell Technologies) at 37 °C and 5% CO_2_. Cultures were fed daily. For passaging of iPSC cultures, differentiated colonies were manually scratched off the bottom of the well and media was aspirated. Fresh media was added and passaged 1:6 using the StemPro EZpassage tool (Life Technologies) as per manufactures instructions.

### NPC derivation

iPSCs were cultured with the addition of 10 ng/mL StemBeads FGF-2 (StemCultures). Upon reaching 80 % confluency, the media was changed to mTeSR™-E8™ supplemented with 10 ng/mL StemBeads FGF-2 and 10 μM Y-27632 (Sigma-Aldrich). After 24 h, cells were washed with PBS, dissociated with gentle cell dissociation reagent (StemCell Technologies) and incubated for 10 min (37 °C, 5 % CO_2_). Cells were gently pipetted up and down to ensure all cells had dislodged and formed a single cell suspension. Cells were then centrifuged (300 g, 5 min) and plated at a density of 90,000 cells/mL in STEMdiff Neural Induction Medium (StemCell Technologies) with SMADi supplement (StemCell Technologies) (NIM) and 10 μM Y-27632 on an ultra-low attachment U-bottom 96-well plate (100 μL/well) (Costar) and incubated at 37 °C, 5 % CO_2_. ¾ of the media was changed daily for 4 days, ensuring not to remove the embryoid bodies. Embryoid bodies were aspirated using a 200 μL wide-bore pipette tip and plated onto Matrigel-coated (0.08 mg/well) 6-well plates (10-13 embryoid bodies/well) in NIM. Media was fully replaced daily for 7 days. Cells were washed with DMEM/F12 and incubated for 70 min (37 °C, 5 % CO_2_) in STEMdiff Neural Rosette Selection Reagent (StemCell Technologies). Rosettes were lifted by gently dispensing DMEM/F12 onto the colonies, centrifuged (350 g, 5 min), plated onto a Matrigel-coated (0.08 mg/well) 6-well plate in NIM and incubated for 24 h (37 °C, 5 % CO_2_). Media was switched to neural progenitor cell (NPC) media containing DMEM/F12, 1 x N2 (Invitrogen), 1 x B27-RA (Invitrogen) and 20 ng/ml FGF2 (Abcam) and changed daily. NPCs were passaged after 4 days using accutase (Sigma-Aldrich) (5 min, 37 °C, 5% CO_2_) and maintained on Matrigel-coated (0.08 mg/well) 6-well plates. Further passaging was done roughly 1:3 every week.

### Astrocyte derivation

NPCs were plated at 15,000 cells/cm^2^ on Matrigel-coated (0.08 mg/well) 6-well plates in NPC medium and incubated for 24 h (37 °C, 5 % CO_2_). Media was switched to astrocyte medium (astrocyte basal medium (ScienCell), 2 % FBS, astrocyte growth supplement and 10 U/mL penicillin/streptomycin solution) and were fed every other day. Cells were passaged with accutase (5 min, 37 °C, 5 % CO_2_) when reaching 90 % confluency, centrifuged (300 g, 5 min) and plated at the original plating density. After 30 days in astrocyte medium, astrocyte identity was then validated using IHC and used for experiments.

### iBMEC Derivation

The protocol for derivation of iPSC-derived BMECs follows that outlined in Neal et al. (2019). Briefly, iPSCs were singularised with accutase, resuspended in mTeSR1 with 10 μM Y-27632 (In Vitro Technologies) and seeded at 15,800 cells/cm^2^ on 16.6 μg/cm^2^ Matrigel-coated plates. 24 h later, medium was changed to E6 medium (A1516401), and changed daily for 4 days. Medium was then changed to human endothelial serum-free media (hESFM) with 0.5 % B27 (Life Technologies), 20 ng/mL bFGF and 10 μM all-trans retinoic acid (RA; Sigma-Aldrich). After 48 h cells were then singularised and plated onto substrates coated with 400 μg/mL collagen IV (Sigma-Aldrich) and 100 μg/mL fibronectin (F1141). 24 h later, RA and bFGF were removed from the media. Assays were performed 24 h after the removal of bFGF and RA.

### Neuron Derivation

Neuron derivation followed a previously published protocol outlined in (Bardy et al., 2015). For differentiation of neurons in 2D, NPCs were plated on Matrigel-coated (0.08 mg/mL) 96-well plates at 40,000 cells/well in 100 μL NPC media. The following day 50 μL of neuron media (BrainPhys media supplemented with 1:50 NeuroCult SM1 supplement, 1:100 N2 Supplement-A, 20 ng/mL BDNF, 20 ng/mL GDNF, 1 mM cyclic-AMP, 200 nM vitamin C) (StemCell Technologies) was added to the wells. A half media change was conducted every 2-3 days for 4 weeks. For 3D differentiation, NPCs were printed using the RASTRUM 3D Bioprinter at a final concentration of 5 x 10^6^ cells/mL of matrix. A full media change of NPC media was conducted every day for 3 days. 4 days post-printing, 50 μL of neuron media was added to the wells and a half media change was conducted every 2-3 days for 4 weeks.

